# The prospective sense of agency is rooted in local and global properties of intrinsic functional brain networks

**DOI:** 10.1101/2020.02.14.948885

**Authors:** Simone Di Plinio, Mauro Gianni Perrucci, Sjoerd J.H. Ebisch

## Abstract

The sense of agency (SoA) refers to a constitutional aspect of the self describing the extent to which individuals feel in control over their actions and consequences thereof. Although the SoA has been associated with mental health and well-being, it is still unknown how interindividual variability in the SoA is embedded in the intrinsic brain organization. We hypothesized that the prospective component of an implicit SoA is associated with brain networks related to SoA and sensorimotor predictions on multiple spatial scales. We replicated previous findings by showing a significant prospective SoA as indicated by intentional binding effects. Then, using task-free functional magnetic resonance imaging (fMRI) and graph analysis, we analyzed associations between intentional binding effects and the intrinsic brain organization at regional, modular, and whole-brain scales. The results showed that inter-modular connections of a fronto-parietal module including the premotor cortex, supramarginal gyrus, and dorsal precuneus are associated with individual differences in prospective intentional binding. Notably, prospective intentional binding effects were also related to global brain modularity within a specific structural resolution range. These findings suggest that an implicit SoA generated through sensorimotor predictions relies on the intrinsic organization of the brain connectome on both local and global scales.

## 1. INTRODUCTION

The sense of agency (SoA) concerns the experience of oneself as the source of one’s actions and their consequences. This phenomenon is often investigated as a state- or event-related subjective experience from which researchers can extrapolate the extent to which an individual feels in control over his or her behavior, thoughts, and the environment. Acquiring a sense of agency is considered a fundamental step in cognitive development (Ruvolo et al., 2015) as well as in human evolutionary adaptation (Taylor et al., 2014). The experience of a lower sense of agency can impair behavioral performances on cognitive tasks (Schooler et al., 2014), cause a loss of awareness (Berberian et al., 2017), and negatively impact the quality of everyday life (Bandura, 2006; Renes and Aarts, 2017) and mental health (Moore and Fletcher, 2012; de Bézenac et al., 2018).

The implicit sense of agency can be measured by intentional binding, that is, the shift of the subjectively perceived timing of a performed action toward the consequences of that action (Haggard et al., 2002; Moore and Obhi, 2012). Intentional binding can rely on two mechanisms (Moore and Haggard, 2008): firstly, prospective intentional binding is based on the probabilistic and context-dependent coding of action consequences supported by internal predictive models; secondly, outcome-dependent, a posteriori inferences about action consequences support retrospective intentional binding.

Studies on intentional binding have highlighted many substantial interindividual differences in the sense of agency and their implications for mental health. For instance, while healthy individuals show a significant prospective component of the sense of agency as measured through intentional binding (Moore and Haggard, 2008), increased schizotypal traits and schizophrenia have been associated with weakened prospective and increased retrospective intentional binding effects (Moore et al., 2011). It has also been shown that psychosis-related and positive social personality traits may predict decreases or increases in the prospective intentional binding depending on individual environmental control (Di Plinio et al., 2019a). Furthermore, intentional binding differs between high- and low-hypnotizable individuals (Lush et al., 2019) and correlates with narcissistic personality traits (Hascalovitz and Sukhvinder, 2015). These studies consistently demonstrate interindividual differences in the tendency to experience an enhanced or reduced sense of agency.

Increasing evidence from analyses based on graph theory (Bullmore and Sporns, 2009; Rubinov and Sporns, 2010) suggests that the intrinsic organization of brain networks contributes to the predisposition of an individual’s typical behavioral patterns (Gallen and D’Esposito, 2019). Intriguingly, many brain regions implicated in agency, including the premotor cortex, the inferior parietal lobule, the anterior insula, the cerebellum, and the precuneus (Farrer et al., 2003; Desmurget et al., 2009; Chambon et al., 2013; Rae et al., 2014; Chambon et al., 2015; Króliczak et al., 2016; Haggard, 2017), have been described as information hubs in the human brain (Buckner et al., 2009; Power et al., 2013). Hence, these regions could act as local multimodal integrators of sensorimotor information (van den Heuvel and Sporns, 2013) within motor control networks to support the sense of agency.

Besides, the potential link between agency and the intrinsic brain architecture may concern not only local elements of the brain (i.e., specific regions/nodes or networks/modules) but also whole-brain parameters (Mišic and Sporns, 2016). For example, it has been proposed that complex neurocognitive processes may also arise from the brain’s global modular structure, that is, to the degree of integration and segregation among brain subsystems (Ito et al., 2019). This view is supported by recent studies that linked the brain’s intrinsic organization to the predisposition of individual behavioral patterns (Godwin et al., 2015; Hilger et al., 2017; Gupta et al., 2018). Specifically, the prospective sense of agency reflects a constitutional aspect of self-awareness (Gallagher, 2000; Prinz, 2012) that emerges from sensorimotor prediction models (Wolpert et al., 2001; Clark, 2013; Haggard, 2017). Predictive coding, putatively grounded in the integration of actions and their outcomes (Kilner et al., 2007), is a general property of the brain that is fundamental for both its development (Wolpert, 1997) and its adaptability (Sato and Yasuda, 2005). Indeed, better contextual predictions mean smaller prediction errors (Friston and Kiebel, 2009) and increased adaptability of the brain to external demands.

Whether the intrinsic functional connectivity patterns of regions associated with the sense of agency and sensorimotor predictions predispose individual propensities regarding the sense of agency is still unknown. Given these studies, the quality of these mechanisms likely depends on both local and global features of the brain system, whereas a high efficiency of the brain in generating accurate predictive schemes may be necessary for a healthy sense of agency.

The purpose of this functional magnetic resonance imaging (fMRI) study was to investigate if and how an implicit sense of agency, reflected by intentional binding, is associated with the brain’s intrinsic modular organization. To that end, we first assessed the prospective and retrospective components of the sense of agency following previous studies (Haggard et al., 2002; Voss et al., 2010; Di Plinio et al., 2019a) in a sample of healthy individuals (N = 39). Then, we studied the associations between implicit measures of the sense of agency (prospective and retrospective intentional binding) and intrinsic functional network features during a task-free (i.e., resting) state on three different, complementary spatial scales: nodal (brain regions), modular (brain networks), and global (whole brain).

We hypothesized that the neural processing of the individual sense of agency may be specifically modulated according to individual patterns of intrinsic brain connectivity. Given that predictive mechanisms are more likely to be embedded in the intrinsic brain organization (Körding et al., 2007; Kannape and Blanke, 2012; Apps and Tsakiris, 2014), while retrospective mechanisms are more likely related to task-evoked signals, we mainly expected that the prospective sense of agency could be related with task-free fMRI measures. On the one hand, we supposed that the segregation of functional brain subsystems (i.e., modules) as indexed by global modularity may favor an efficient general organization of information processing in the brain, allowing higher adaptability of sensorimotor predictions and, consequently, a higher sense of agency indexed by the prospective component. On the other hand, we also hypothesized that topological features of specific regions or subsystems of the brain may be involved in the transfer and integration of sensorimotor information across brain subsystems/networks. Specifically, on the nodal and modular level, we expected that prospective intentional binding may be associated with enhanced nodal efficiency (van den Heuvel and Sporns, 2013) or participation coefficients (Guimerà and Nunes Ameral, 2005) for regions such as the inferior parietal lobule, the precuneus, the premotor cortex, the insula, and the cerebellum.

## 2. MATERIALS and METHODS

### 2.1. Participants

Thirty-nine healthy Italian adults (19 females and 20 males, aged 23 ± 2; 35 right-handed and 4 left-handed) without a history of psychiatric or neurological disease and contraindications for MRI scanning participated in the experiment. The experiment was approved by the local ethics committee. All participants had a normal or corrected-to-normal vision and provided written informed consent before taking part in the study in accordance with the Declaration of Helsinki (2013). Each participant performed the behavioral sense of agency task one to three days before the MRI acquisition.

### 2.2. Behavioral procedure

All participants performed a well-established sense of agency paradigm (Haggard et al., 2002; Voss et al, 2010; Di Plinio et al., 2019a). On each trial, the participants performed voluntary, self-initiated keypresses with the right index finger, while watching a clock hand rotating on a screen. The clock was labelled using a circular scale with numbers positioned at 5-minutes intervals. The participants’ task was to judge the timing of the keypress, i.e., the clock time indicated by the clock hand when they pressed the key. The participants started each trial by voluntarily pressing a button with the left index finger. The clock hand rotation began at a random position in each trial and performed a full rotation every 2560 ms. The inter-trial interval was jittered between two and three seconds.

Two experimental conditions of action timing were used in the sense of agency task. In the 50% probability condition, participants heard a tone following the keypress in 50% of the trials. Instead, in the 75% probability condition, the keypress was randomly followed by a tone presentation in 75% of the trials. The tone consisted of a single pulse at 1,000 Hz and lasting 200 ms. The tone was presented binaurally 250 ms after the keypress. The task was performed using the E-Prime software (Schneider et al., 2012), while participants were sitting approximately 60 cm from the 20-inches monitor. Two blocks for each condition (50%, 75%) of the sense of agency task were performed by each participant. Each block consisted of 32 trials. The participants also performed a fifth, baseline block composed of 32 trials in which no tones occurred. Block order was pseudo-randomized across participants. The whole procedure lasted approximately 45 minutes for each participant.

The variable measured in each trial of the sense of agency task was the perceptual shift (Δ). The perceptual shift is defined as the time difference between the real keypress and the perceived keypress indicated by the subject in each trial, and it is expressed in milliseconds. To note, although the perceptual shift and the phenomenon of the intentional binding are related, they should not be confused: while the former represents the divergence between real and perceived keypress, the second is a phenomenon that concerns the shift of the perceived timing of action toward the action consequences (sound).

The participants were instructed (i) to avoid the planning of their keypresses, (ii) to avoid answering in a stereotyped way, and (iii) to avoid answering before the end of the first full clock rotation. A brief training session was performed by each participant before the experiment. The task is illustrated in Figure 1.

**Figure 1.**
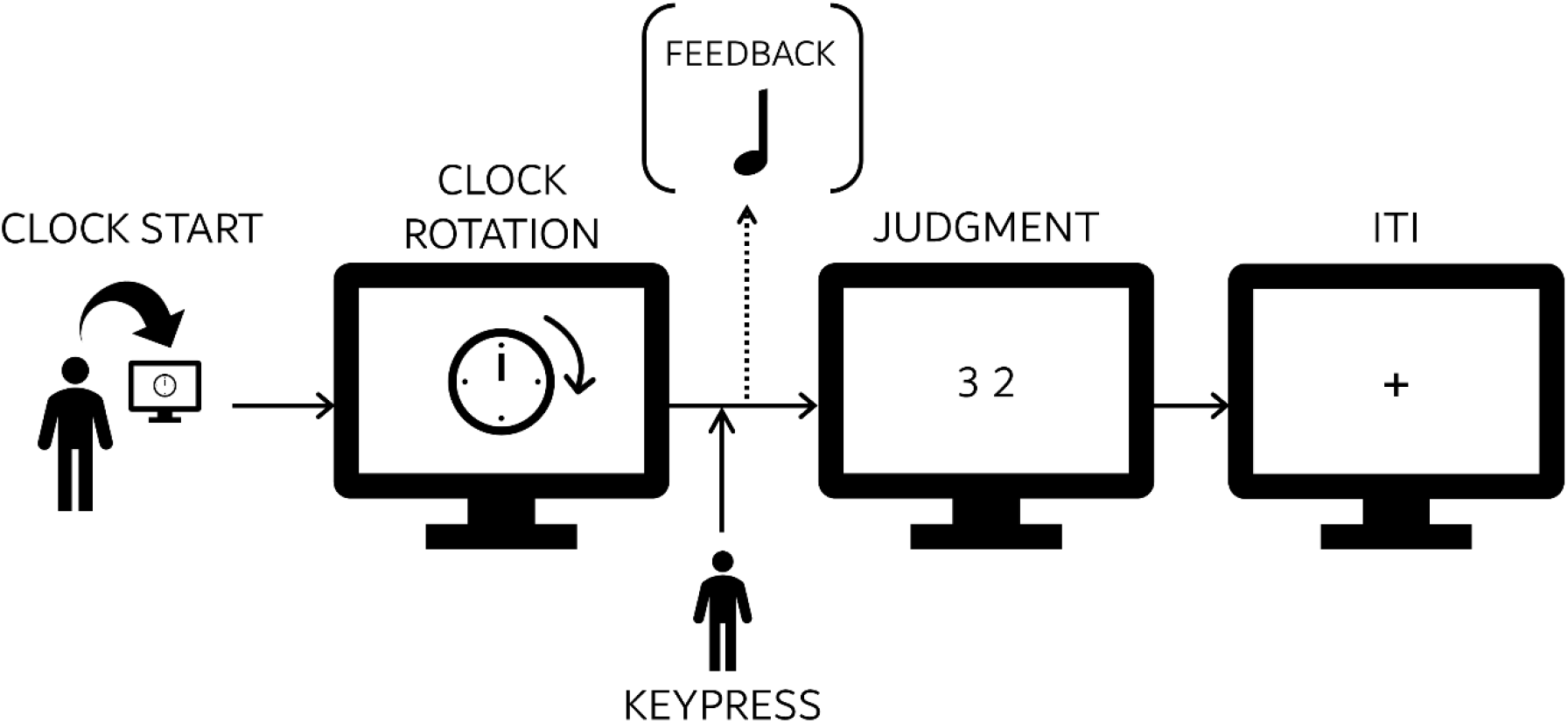
Behavioral task. Participants performed voluntary, self-paced keypresses with the right index finger while watching a clock hand rotating on a screen. Participants started each trial by themselves pressing a button with the left index finger, and they were instructed to judge the clock ‘time’ indicated by the clock hand when they pressed the key. After the keypress, depending on the trial and on the experimental condition, auditory feedback (or nothing, contingent on the current trial) was binaurally presented through headphones.

### 2.3. Data acquisition

Each participant performed two consecutive task-free fMRI runs, each consisting of 376 volumes. The participants were instructed to watch a white fixation cross on a black screen without performing a cognitive task. Each run lasted approximately 7.5 minutes. Functional images were acquired using a Philips Achieva 3T scanner installed at the Institute for Advanced Biomedical Technologies (Gabriele d’Annunzio University, Chieti-Pescara, Italy). Whole-brain functional images were acquired with a gradient echo-planar sequence using the following parameters: repetition time (TR) = 1.2 s, echo time (TE) = 30 ms, field of view = 240×240×142.5 mm, flip angle = 65°, in-plane voxel size = 2.5 mm^2^, slice thickness = 2.5 mm. A high-resolution T1-weighted whole-brain image was also acquired after functional sessions using the following parameters: TR = 8 ms, TE = 3.7, FoV = 256×256×180 mm, flip angle = 8°, in-plane voxel size = 1 mm^2^, slice thickness = 1 mm.

### 2.4. Preprocessing of MRI data

The preprocessing steps and the analysis of functional images were implemented in AFNI (Cox, 1996). A slice-timing correction was applied to remove differences in acquisition times between the slices. The obtained functional images were deobliqued, despiked, corrected for time-shifted acquisition, and motion-corrected using a six-parameter rigid body realignment before being aligned to the Montreal Neurological Institute (MNI) standard brain using non-linear warping. During the preprocessing steps, motion parameters for each participant were collected. The functional images were scaled to have voxels with an average value of 100 to translate the unitless BOLD signal acquired by scanners to the “BOLD percent signal change” as a more interpretable index (Chen et al., 2017). A Gaussian filter of 5 mm full-width at half-maximum was applied to spatially smooth the functional images.

Finally, the time series were censored (volumes with 10% or more outliers across voxels and volumes with Euclidean norm of the motion derivative exceeding 0.2 mm were excluded as suggested in Power et al., 2014), and band-pass filtered (frequency interval: 0.01-0.10 Hz) in a single regression step (Caballero-Gaudes and Reynolds, 2017) in which the motion parameters were also included as noise regressors together with white matter and cerebrospinal fluid signals. The mean framewise displacement was also added as an additional covariate of no interest (Power et al., 2014; Ciric et al., 2018). We did not regress out the global signal because it is a controversial approach (Saad et al., 2012) and potentially introduces spurious negative correlations (Weissenbacher et al., 2009).

### 2.5. Behavioral data analysis

The average baseline perceptual shift (Δ_B_) was subtracted from the average perceptual shift in each experimental condition and each participant (Voss et al., 2010; Di Plinio et al., 2019a). Linear mixed-effects analyses were implemented using *tone probability* (levels: 50% and 75%) and *trial type* (levels: Action & tone, action only) as fixed effects, while a random intercept was added at the subject level. The dependent variable was the average perceptual shift minus the baseline perceptual shift (Δ-Δ_B_). To identify the significance of prospective/retrospective components, we performed multiple comparisons using Tukey’s honest significant difference test. The homogeneity of residuals was assessed using the D’Agostino-Pearson test.

We also implemented a bootstrap procedure to estimate the distributions of the effect sizes and to investigate the modulation of the prospective and retrospective components (Kirby and Gerlanc, 2013). The prospective component was estimated by contrasting the perceptual shifts during ‘action only’ trials in the 75% probability condition versus perceptual shifts during ‘action only’ trials in the 50% probability condition, thus, reflecting the contextual effect of tone probability on perception. The retrospective component was estimated by contrasting the perceptual shifts between ‘action+tone’ and ‘action only’ trials within the 50% probability condition, thus, reflecting the outcome-dependent effect on perception. The effect size (Hedge’s g) was estimated in 10,000 bootstrap cycles with replacement for both the prospective and retrospective components.

### 2.6. Connectomics

The two task-free runs were concatenated and whole-brain functional connectomes were formed using a set of 418 nodes derived from the cortical (346 parcels) and subcortical (40 grey nuclei) atlases from Joliot and colleagues (2015) plus the cerebellar (32 nodes) atlas from Diedrichsen and colleagues (2009). Functional connectivity was calculated using the z-Fisher transform of the Pearson correlation among average time series extracted from the voxels within each node. Connectomes were used to build binary undirected graphs after thresholding (the 10% strongest connections were retained). Graph analyses were performed within Matlab using the Brain Connectivity Toolbox (Rubinov and Sporns, 2010). Modular subdivisions of the brain were visualized using the software BrainNet Viewer (https://www.nitrc.org/projects/bnv/; Xia et al., 2013) implemented in MatLab. Formal definitions of the metrics used in graph analysis are reported in Table I.

To find the optimal community structure of the networks, we implemented modularity maximization (Porter et al., 2009) through the application of the widely used and robust Louvain algorithm (Blondel et al., 2008; Lancichinetti and Fortunato, 2009). An iterative fine-tuning process was used to find subject-specific optimal modular structures (Sun et al., 2009) and to handle the stochastic nature of the Louvain algorithm (Bassett et al., 2011). In the first step, each node was assigned to a distinct module. Then, the optimal community was detected using the Louvain algorithm and the modularity Q was estimated. This procedure was repeated using the optimal community found in the previous cycle as the starting community affiliation vector, and the process was repeated until Q could not be increased anymore. The agreement matrix was calculated as the matrix whose elements represented the number of times two nodes were assigned to the same module across subjects. The group-level modular structure was achieved from the agreement matrix using a community detection algorithm developed for the analysis of complex networks (Lancichinetti and Fortunato, 2012), with the number of repetitions set to 1,000.

During the detection of optimal communities of nodes in the brain, the structural resolution parameter (γ) plays an important role since it weights the null model in the modularity estimation. In the present study, γ was varied in the interval [0.3-5.0] with steps of 0.1 to avoid biasing the subsequent analyses (Betzel et al., 2016). Then, the similarity between the consensus structure and the community structure across participants was calculated using the adjusted Rand coefficient (Traud et al., 2011) for each γ. Finally, an automated maximization algorithm was implemented in Matlab to detect one or more γMAX associated with local maxima of the Rand coefficient among the 48 tested gammas. The Newman-Grivan procedure was employed to detect significant modules in the consensus structure(s) (Newman and Girvan, 2004), as follows. Null models (10,000) were generated using a random permutation of module assignments while maintaining the original size and number of modules. Within each null model, the modularity contribution QC of each module was calculated as the summation of the real minus the expected number of connections between node pairs. The expected number of connections between each node pair was weighted with node degree (Newman and Girvan, 2004; Betzel et al., 2016). Only modules in which the real modularity contribution was greater than 99.9% of the simulated modularity contributions (p < .001) were considered significant.

The percentage of each predefined network (Di Plinio and Ebisch, 2018) covered by each module was estimated to characterize the modules’ functional fingerprints.

### 2.7. Brain-behavior relationship analysis

Global statistics of modularity (Q) and global efficiency as well as nodal statistics of participation (Guimerà and Nunes Ameral, 2005) and efficiency (van den Heuvel and Sporns, 2013) were extracted and analyzed concerning the prospective and retrospective component of the sense of agency.

A two-step robust weighted regression was applied to analyze the participation coefficients excluding poor fitting and overfitting. In the first step, the model was fitted using all the module’s nodes as levels of a random grouping variable, and the prospective component was used as a continuous predictor. In the second step, the analyses were repeated after removing nodes violating model assumptions, that is, with inhomogeneous residuals within the corresponding random grouping as indicated by Anderson-Darling normality tests. Regressions were performed separately for each module to detect module-specific associations between participation measures and the sense of agency. Results were corrected for multiple comparisons after model diagnostics and outlier removal. Best linear unbiased predictors (BLUPs) were extracted to estimate the effect at the nodal level and highlight nodes with the highest contributions whereas significant effects were detected (Liu et al., 2008). Individual Conditional Expectation (ICE) plots were generated to visualize significant effects at both the nodal and modular level (Goldstein et al., 2013). The impact of significant results was tested also considering different graph density thresholds (10, 15, 20, 25, 30%) through the cost integration approach (Ginestet et al., 2011; Hilger et al., 2017), conforming to current neuroscientific standards (Nichols et al., 2017; Van den Heuvel et al., 2017). A Bayesian bootstrap procedure was employed to estimate the reliability of the beta coefficients from the regression analyses (Rubin, 1981).

Starting from the hypothesis that the individual intrinsic brain signatures may be associated with the implicit sense of agency depending on the structural resolution of the modular architecture, and in line with recent multi-resolution approaches (Jeub et al., 2018; Chen et al., 2018), the modularity Q was analyzed using all the γ values in the interval [0.3-5.0]. Three different analyses were implemented to test these associations. Firstly, a simple linear model was fitted for each value of γ using the modularity Q as the dependent variable and the prospective component as a continuous predictor. Secondly, the link between modularity and the sense of agency was tested in a multivariate model including all the structural resolutions in a single analysis, with levels of γ being used as repeated measures. Thirdly, a penalized regression (elastic net, Zou and Hastie, 2005) was adopted to account for the many potentially correlated variables, i.e., modularity values at different levels of γ. The optimal regularization coefficient of shrinkage (λ) for the penalized regression was chosen as the λ minimizing the mean squared error in 100 Monte Carlo cycles of two-folded cross-validations (Hastie, et al., 2009). Since the LASSO-penalty in the regression constraints irrelevant variables (irrelevant γs) to be zero, the stability of feature selection was estimated by calculating the frequency of non-zero coefficients in 500 bootstrap cycles (threshold: p < .05). It can be argued that functional connectivity may depend on the level of musical expertise, the fitness of the individuals, handedness, or sex (Herting and Nagel, 2013; Kühnis et al., 2014; van der Westhuizen et al., 2017). To this aim, the analyses described above were also performed adding years of sports training and music expertise as covariates. Additionally, the interaction of the measures of agency and participant’s sex and handedness was considered, and we repeated all the analyses using weighted graphs (Sporns and Betzel, 2016) to explore the stability of the results in relation to the methodology used.

## 3. RESULTS

### 3.1. Behavioral Results: confirming a significant prospective sense of agency in healthy subjects

The average perceptual shifts are reported in Table II and illustrated in Figure 2a. As expected, we found a strong prospective component, while the retrospective component was nearly zero. This effect was confirmed by the results of the mixed-effects model. Significant effects were found for the factor *tone probability* (p = .02, F_(1,160)_ = 5.7) and for the interaction between *tone probability* and *trial type* (p = .01, F_(1,160)_ = 6.1). The factor *trial type* by itself was not significant (p = .38, F_(1,160)_ = 0.8). The statistics observed for the intercept (p = .06, F_(1,160)_ = 3.4) showed that the average perceptual shift (Δ) in the experimental conditions was different from the baseline perceptual shift (Δ_B_), though only a trend was observed. The prospective component was confirmed by contrasting perceptual shifts between the ‘action only’ trials of the two probability conditions (p < .001, Tukey’s HSD test).

**Figure 2.**
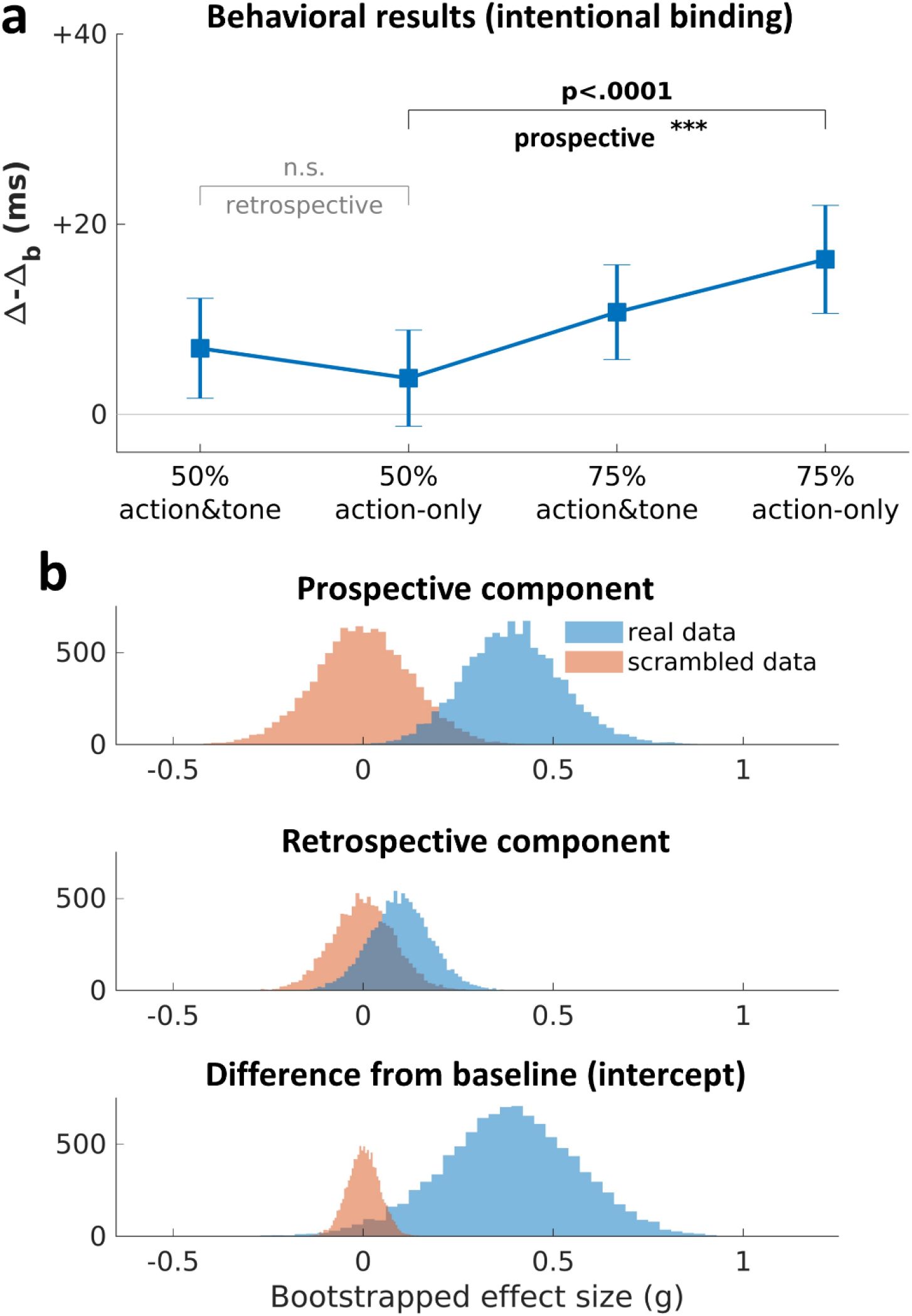
Behavioral results. **(a)** Average values and standard errors of perceptual shifts relative to the baseline condition (Δ-ΔB), which represent measures of intentional binding. As expected, participants showed a significant effect relative to the prospective component. **(b)** Bootstrapped effect sizes (Hedges’ g) for the prospective component, retrospective component, and intercept. Null distributions were created scrambling data between experimental conditions in 10,000 bootstrap cycles. While a medium-to-large effect was observed for the prospective component (upper panel) and the intentional binding (lower panel), the retrospective component showed a close to zero effect size (middle panel). These results replicate findings from Di Plinio et al. (2019a).

The results of the bootstrap procedure to estimate the effect sizes are shown in Figure 2b and indicated a medium-to-large effect size regarding the prospective component (average Hedges’ *g* = 0.39, bootstrapped 95% CI: [0.19 ~ 0.61]). Effect sizes related to the retrospective component were rather irrelevant (*g* = 0.10, 95% CI: [-0.04 ~ 0.23]). Finally, the difference from the baseline was medium-to-large (*g* = 0.38, 95% CI: [0.08 ~ 0.67]).

### 3.2. Graph Analyses: the modular brain structure

Three resolution parameters, corresponding to γ_1_=0.6, γ_2_=1.2, and γ_3_=1.9 were found to maximize the adjusted Rand coefficients (Figure 3a). We report results related to the modular structure associated with γ_3_ since significant relationships with the sense of agency were found within this modular configuration. Group average matrices of functional connectivity are reported in Figure 3b and show the nine significant modules found with γ_3_ (M1-9, p < .001 using the Newman-Girvan procedure).

**Figure 3.**
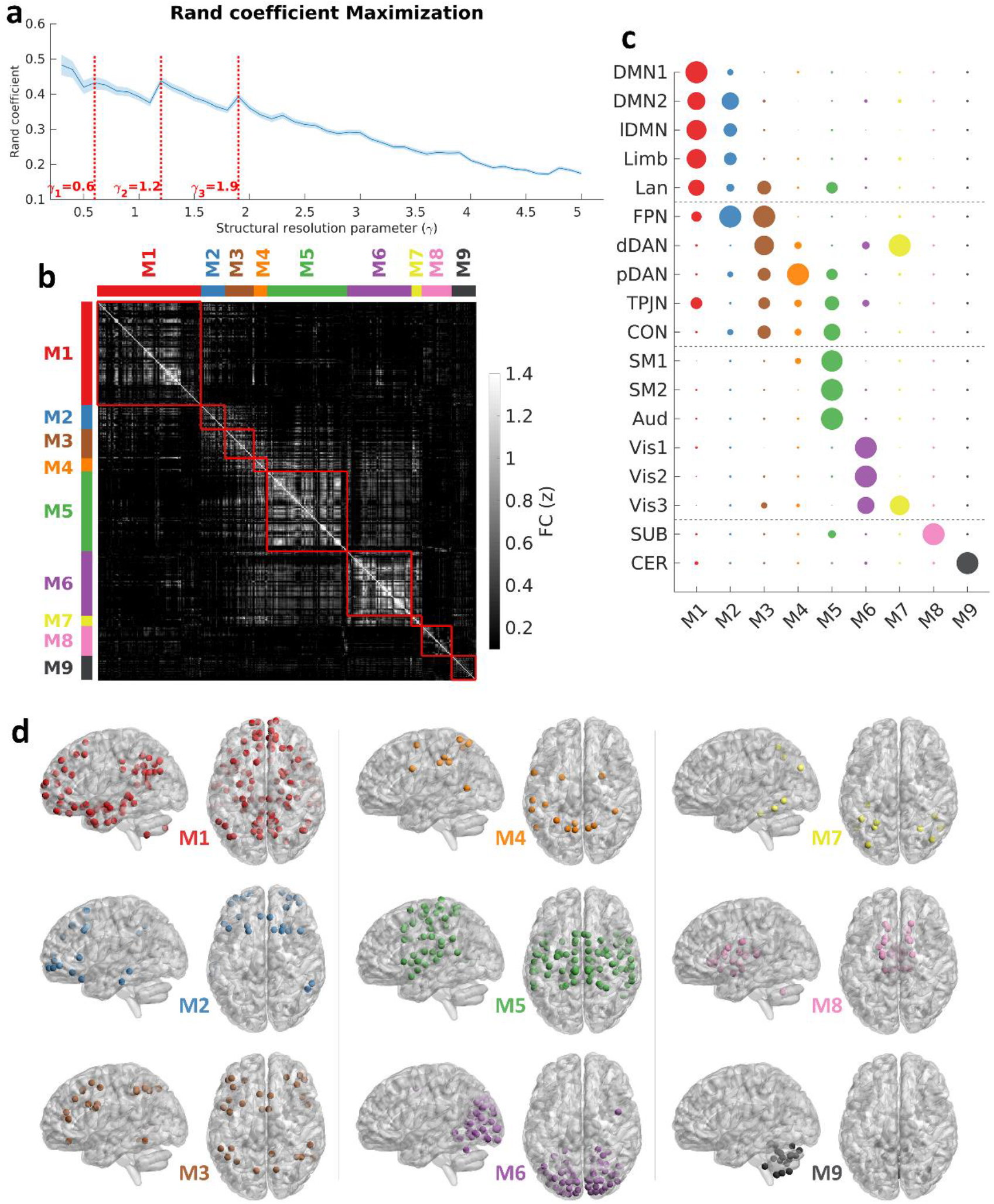
Consensus modularity. **(a)** Local maxima of the Rand coefficient were detected for three gamma values: γ_1_ = 0.6; γ_2_ = 1.2; γ_3_ = 1.9. **(b)** Group average functional connectivity matrix (γ_3_ = 2.1). Each significant module (p < .001, comparison with 10,000 null models following the Newman-Girvan procedure) is enclosed in a square, distinguishing within-modules (inside the squares) and between-modules connections (outside the squares). **(c)** Functional fingerprints of the nine modules identified with γ_3_ = 1.9 as compared with predefined brain networks (Di Plinio and Ebisch, 2018). The width of each circle represents the amount of overlap between each network and each module, calculated as the normalized percentage of each network covered by each module. **(d)** Modular structural topographies with γ_3_ = 1.9. A full list of regions included in each module is available upon request to the corresponding author.

Anatomical fingerprints of the nine modules with respect to predefined brain functional networks (Di Plinio and Ebisch, 2018) are depicted in Figure 3c. The consensus community detection (with γ3) detected five modules encompassing associative cortices in the frontal, parietal, and temporal lobes (M1-4, M7). Among these modules, M1 was mainly related to default mode and language systems, M2 incorporated frontal, insular, and cingulate regions often ascribed to salience and control networks, M3 encompassed high-order executive and attentional regions, M4 included regions associated with motor attention and social interactions, and M7 included regions associated with late stages of visual processing. Furthermore, two sensory modules were detected, M5 and M6, that were related to sensorimotor/auditory and visual networks, respectively. Among the last two modules, M8 included hippocampal and amygdala regions, while M9 included cerebellar structures. Module topographies are illustrated in Figure 3d.

### 3.3. Modular and nodal associations with a prospective sense of agency

A significant association between the participation coefficient and the prospective component of the sense of agency was found at the modular level with γ_3_. The network participation of M4 (motor attention module) was positively associated with the prospective component of the sense of agency (t_(544)_ = 3.3, p = .001, β = 3.6e-4 CI: [1.4e-4 ~ 5.7e-4]; adjusted R^2^ = 0.78). Figure 4a represents the nodal ICE plot and shows the effect both at the modular (thick line) and nodal level (shaded lines). The nodes that showed stronger effects are represented by larger spheres with a higher color intensity in Figure 4b. Scatter plots and effect sizes (β_s_) for the nodes in M4 showing the largest effects are shown in Figure 4c. These nodes were the left precentral gyrus, postcentral gyrus, superior parietal cortex, and supramarginal gyrus (average t = 4.8, p < .001), which are reported in Figure 4c. The significant results were quite consistent and remained significant also using bootstrap procedures and using different graph density thresholds following the cost integration approach (Figures 4d-e). Post-hoc analyses using different gamma values revealed that this association also occurred with other gamma values in which a fronto-parietal motor-attention module was detected (specifically, when γ was equal to 2.5, 4.0, and 4.3). Measures of nodal efficiency were not significantly associated with intentional binding.

**Figure 4.**
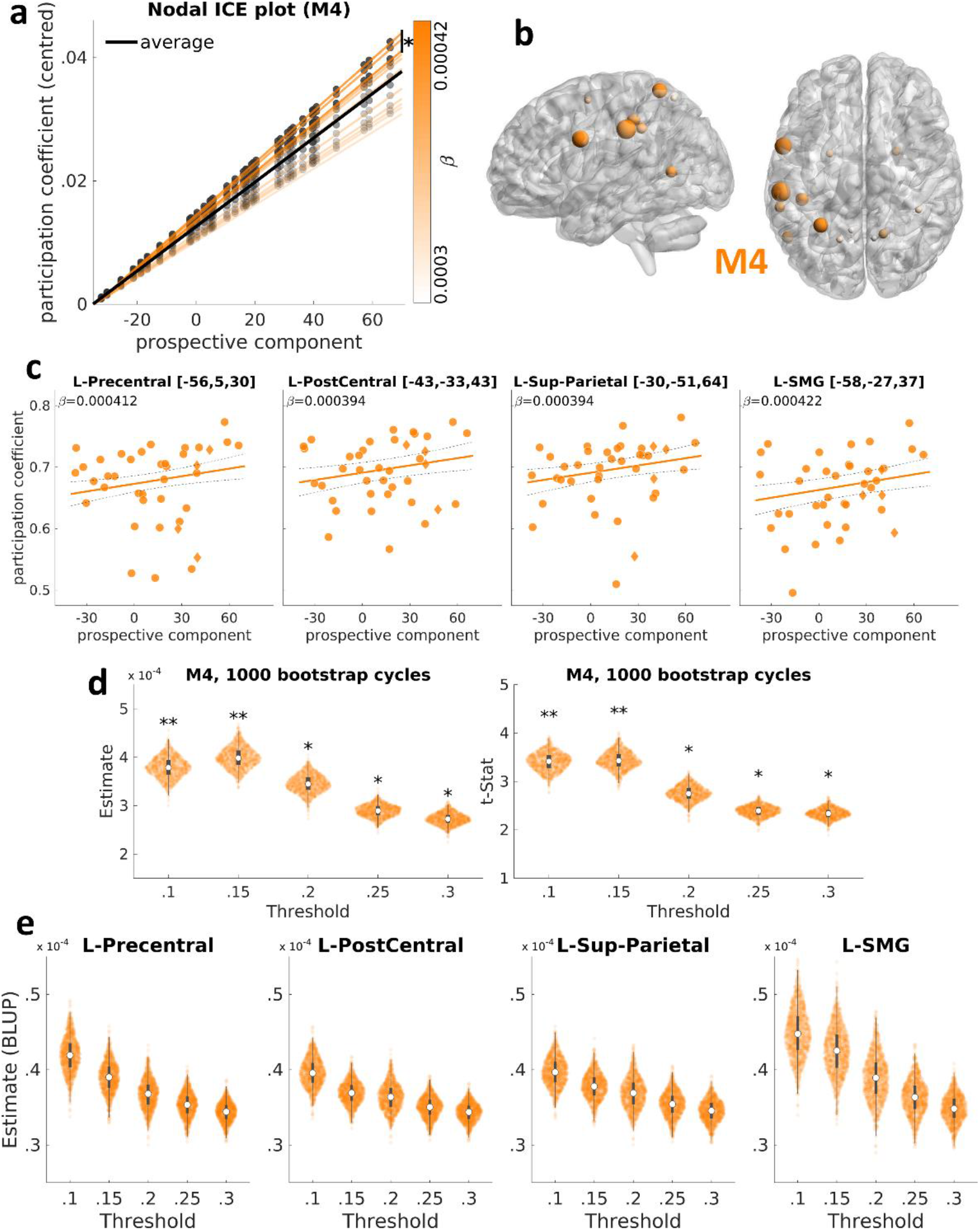
Associations between agency and nodal/modular participation. **(a)** Individual Conditional Expectation (ICE) plot for the module M4 which illustrates the relationships between the prospective component and network dmeasures of participation. Each slope represents the zero-centered predictions of a node’s participation coefficient (Y-axis) for a given individual score in the prospective component (X-axis). Color intensity indicates the effect strength (beta value, estimated using Best Linear Unbiased Predictors, BLUPs). The average effect at the module level is indicated by a thick black line (t_(544)_ = 3.3; p = .001; β = 3.6e-4). **(b)** Module M4. The size and the color intensity of the nodes are regulated according to the strength of the associations (β_s_) between the prospective component and participation coefficients. **(c)** Nodes in M4 in which the effect was statistically stronger than the average slope for the module. These nodes were in the left supramarginal gyrus (L-SMG). For each node, the MNI coordinates and the effect sizes are indicated. Diamonds indicate left-handed subjects. **(d)** Estimates and relative standard errors for the bootstrap analysis of interactions between the prospective component of the sense of agency and participation coefficient (module M4) are reported for each graph threshold analyzed (**: p < .005, *: p < .05). Each point represents the beta estimate following a single bootstrap cycle. Box plots are also shown. To note, results were significant using all the threshold range considered (from 0.10 to 0.30). **(e)** Estimates and relative standard errors for the bootstrap analysis of interactions between the prospective component of the sense of agency and participation coefficient (nodal level) are reported for each graph threshold analyzed. To note, results were significant using all the threshold range considered (from 0.10 to 0.30).

Summarizing, since the participation coefficient reflects the extent of nodal interactions with other modules, these results indicate the presence of a fronto-parietal module of which the inter-modular connectivity covaried with the individual implicit sense of agency. Within this module, the strongest contributors to such effect were two nodes located in the left supramarginal and precentral gyri. Covariates included in the analyses, such as sex, handedness, musical expertise indexed by the years involved in musical training, and sportive skills indexed by years of sports training were not significant and did not interact with the effects of the predictive component. No significant effects were found concerning the retrospective component.

### 3.4. The prospective sense of agency is associated with whole-brain modularity

The association between the prospective sense of agency and multi-scale global modularity was investigated in multiple ways. Using linear regression models, significant positive relationships between the two measures were found for γ ranging in the interval [1.3-2.4] (Figure 5a). The t-statistics for the effect of the prospective component on modularity Q with increasing γ are reported in Figure 5b. In particular, the strongest association was found for γ = 1.8 (t_(37)_ = 2.5, p = .01, β = 4.3e-4).

**Figure 5.**
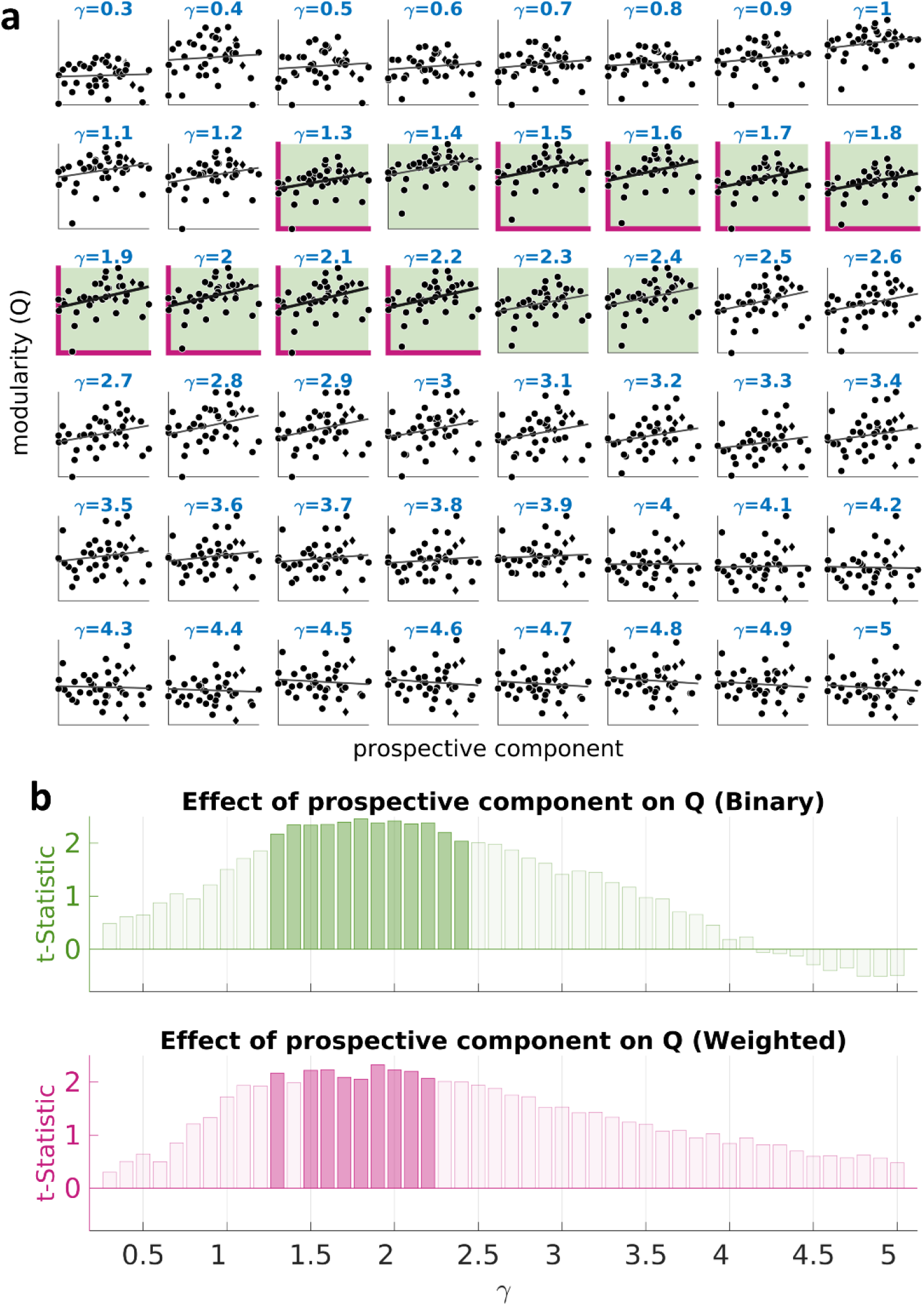
Associations between agency and global modularity: Linear regression. **(a)** Scatter plots representing values of the prospective component (X-axis) against global modularity (Y-axis) for each value of γ tested. Significant (p < .05) interactions in binary graphs are indicated with green background colors. Significant (p < .05) interactions in weighted graphs are indicated with bold purple axes. Diamonds indicate left-handed subjects. **(b)** The t-statistic for the association between global modularity (Q) and the prospective component is represented for each γ and is shown for binary (top panel, green color) and weighted graphs (bottom panel, purple color). Darker green and purple colors significant associations (p < .05).

The same effect was observed using a multivariate model. A higher prospective sense of agency was associated with a higher modularity Q (F_(47)_ = 3.2, p < .001). The effect was still significant after applying the Huynh-Feldt correction for sphericity (p = .049). This relationship was especially strong for γ_s_ in the interval [1.3-2.4]. Figure 6a reports the predicted Q for different values of the prospective component. Although modularity is acknowledged as a measure of the internal cohesion of modules, increasing the structural resolution parameter automatically results in a lower Q. This happens because many connections become cross-modular by using additional smaller modules, that is, more connections are excluded in the estimation of Q. To characterize the relationship between a γ-standardized measure of the modularity and the prospective sense of agency, we also estimated Q_ADJ_, that is, modularity adjusted for the value of γ (Q_ADJ_ = γQ). Figure 6b reports the predicted Q_ADJ_ against increasing values of the prospective sense of agency using the same multivariate model. The association between Q_ADJ_ and the sense of agency was still significant (F_(47)_ = 3.1, p < .001), and remained significant also after Huynh-Feldt correction for sphericity (p = .044).

**Figure 6.**
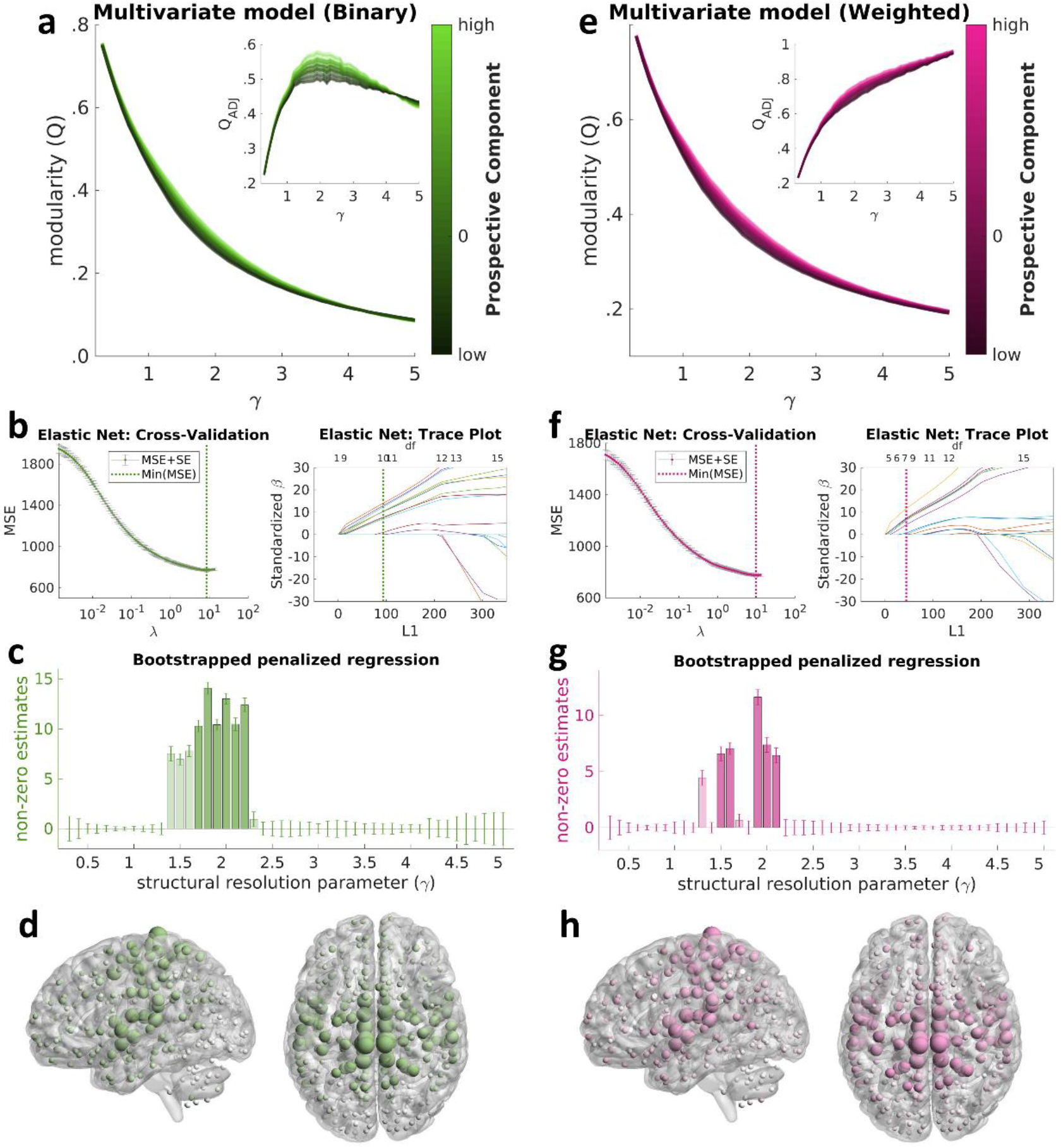
Associations between agency and global modularity: Multivariate and penalized regressions. **(a)** Binary graphs: multivariate model. Predicted values of modularity (Q) and adjusted modularity (Q_ADJ_) with increasing values of γ. The color indicates the value of the prospective component (dark = low prospective component, ≈ −35 ms; green = high prospective component, ≈ 65 ms). **(b)** Binary graphs: penalized regression. Cross-validation for the penalized regression using elastic net (α = 0.5) and the resulting trace plot showing that minimization of the MSE implied a structure with 10 non-zero beta estimates (degrees of freedom: df = 10). **(c)** Binary graphs: penalized regression. Standardized effect sizes associated with each value of γ. The 99% confidence intervals in the figure were obtained through 500 cycles of penalized regression using bootstrap with replacement. Darker green colors indicate gammas associated with significant non-zero betas following the bootstrap procedure (p < .05). Bars indicate 95% confidence intervals. **(d)** Binary graphs: nodal contributions. The size of each node indicates its contribution to the association between global modularity and sense of agency. **(e, f, g, h)** Results of the same analyses for weighted graphs.

The cross-validated mean square errors (MSE) regarding the elastic net penalized model are shown in Figure 6c. The λ associated with the minimum MSE produced a structure of beta estimates with 10 non-zero significant values (Figure 6d). Remarkably, as shown in Figure 6e, the elastic net regression returned non-zero estimates in correspondence to all the significant γ values from the linear regression models. The highest effect was associated with γ = 1.8 (β = 14.1). Modularity values with γ in the interval [1.7-2.2] were significant following the bootstrap procedure used to control for the stability in feature selection (darker colors in Figure 6c).

To further explore the meaning of the results relative to global modularity, we implemented a post-hoc analysis of the association between global modularity and the sense of agency. For each γ and each module MX, we estimated the modularity Q’ excluding the module MX. Then, we linearly regressed Q’ against the prospective component to find the effect size β’. The contribution C of the module MX was estimated as C_MX_ = β – β’, that is, as the difference between the effect size obtained considering the whole brain (β) and the effect size obtained excluding the module (β’). To note, this procedure avoids the introduction of magnitude biases. Finally, values of C_MX_ were associated with each node in MX and summed across γ values. Thus, higher values of C indicate a higher contribution of nodes in the module MX to the association between agency and modularity. The results of this post-hoc analysis are illustrated in Figure 6d and show that nodes in sensorimotor and insular regions were the strongest contributors to the association between global modularity and agency using both binary and weighted graphs. No significant effects were found concerning the retrospective component. Measures of global efficiency were not significantly associated with intentional binding.

Summarizing, the results from linear, multivariate, and penalized regression models showed a significant positive association between the prospective (but not the retrospective) sense of agency and the global modular organization of the brain with medium structural resolutions (with γ in the interval [1.4-2.3]). Importantly, even if it may be considered a sub-optimal model, results from linear regression would allow possible comparisons with studies focalizing on single γ values. Covariates included in the analyses, such as sex, handedness, musical expertise indexed by the years involved in musical training, and sportive skills indexed by years of sports training were not significant and did not interact with the effects of the predictive component.

### 3.5. Results using weighted graphs

The results reported in the previous paragraphs were confirmed using weighted graphs. The modular structure identified with weighted graphs was largely similar to the modular structure obtained through binary graphs, including a default mode module analogous to M1, a salience/control module analogous to M2, a motor attention module analogous to M4 (including supramarginal gyrus, superior parietal cortex, premotor cortex, and dorsal precuneus), a sensorimotor module analogous to M5, a visual module analogous to M6, a limbic module analogous to M8, and a cerebellar module analogous to M9. The network participation of the motor attention module, corresponding to M4, was positively associated with the prospective component of the sense of agency (t_(544)_ = 3.9, p < .001, β =3.1e-4 CI: [1.5e-5 ~ 4.6e-4]; adjusted R^2^ = 0.88), thus, confirming the results from binary graphs. One difference between weighted and binary graphs was related to the contribution of nodes to the effect: in weighted graphs, the strongest effect was associated with nodes in the dorsal precuneus.

The association between the prospective sense of agency and multi-scale global modularity was also confirmed with weighted graphs. Using linear regression models, significant effects were found for γ ranging in the interval [1.4-2.2]. The strongest association was found for γ = 1.9 (t(37) = 2.3, p = .026, β = 4.3e-4). The same effect was observed by using a multivariate model (F_(47)_ = 3.0, p < .001, Figure 6e). The effect was still significant after applying the Huynh-Feldt correction for sphericity (p = .049) and penalized regression, which returned non-zero estimates in correspondence of five of the seven significant γ values from the linear regression models (Figure 6f). Modularity values associated with most γ_s_ in the intervals [1.5-1.6] and [1.9-2.1] were significant following the bootstrap procedure (Figure 6g). The highest effect was associated with γ = 1.9 (β = 11.6). As in the case of binary graphs, the strongest contributors to this association were found in the sensorimotor and insular nodes (Figure 6h)

These results demonstrate an association between the sense of agency and both local and global properties of the resting brain independently of the method used to study the brain connectome (binary graphs, weighted graphs).

## 4. DISCUSSION

We showed that signatures of intrinsic brain activity are associated with the individual sense of agency in healthy adults. We focused on intentional binding as a measure of predictive and retrospective components of an implicit sense of agency (Voss et al., 2010; Haggard, 2017) and on graph-theoretical metrics to describe the intrinsic organization of functional brain networks based on a multi-scale, cross-resolution approach (Bullmore and Sporns, 2009; Rubinov and Sporns, 2010). To the best of our knowledge, this is the first study that traces the experimental measure of the sense of agency to individual differences in the intrinsic functional brain architecture at both local and global scales.

At the regional (nodal) and network (modular) levels, participation coefficients were studied as measures of cross-network integration. Our results revealed an association between the prospective sense of agency and participation coefficients in a fronto-parietal module, labeled M4, which consists of fronto-parietal regions related to motor attention such as the inferior parietal lobule (IPL), premotor cortex (PMC), and dorsal precuneus. Results were significant using both binary and weighted graphs. These same regions have been associated with the sense of control over the environment to predispose the body to motor learning (Wolpert et al., 2011) and to generate internal predictions, which are then compared with perceived sensory stimuli (Gallese, 2000; Desmurget and Sirigu, 2009; Haggard, 2017). For example, IPL is a multimodal integration region (Caspers et al., 2011) involved in the translation of motor intentions into meaningful actions (Farrer et al., 2003; Rae et al., 2014) and in distinguishing the self from other agents to generate a prospective sense of agency (Chambon et al., 2015; Ticini et al., 2018). Furthermore, PMC is involved in monitoring conscious movements (Desmurget et al., 2009) and in higher-order cognitive processes like overt task control (Chambon et al., 2013), while the dorsal precuneus has been associated with both the perceived experience of agency (Cavanna and Trimble, 2006). Accordingly, certain sensorimotor disorders, such as the alien hand syndrome (Assal et al., 2007; Hassan and Josephs, 2016) and impaired movement initiation (Desmurget et al., 2009), have been associated with aberrant functioning of the IPL, and others, such as anosognosia for hemiplegia (Vocat et al., 2019), with impaired functioning of the PMC. These impairments often compromise the online adjustment of motor actions in voluntary actions versus externally driven movements (Fried et al., 2017).

Taken together, these task-related studies have highlighted the role of regions involved in motor attention and motor learning, such as SMG, PMC, and dorsal precuneus, in sensorimotor processes that support the experience of control over actions and their intended consequences. In this context, our results on the local and modular scales make two major contributions. Firstly, behavioral measures of an implicit, prospective sense of agency are related to the cross-module integration of a fronto-parietal subsystem (module M4) encompassing regions related to motor control and sense of agency. Secondly, within this subsystem, the most prominent contribution of inter-network connections to the prospective sense of agency comes from the left PMC, SMG, superior parietal cortex, and postcentral gyrus (using binary graphs) or from bilateral dorsal precuneus (using weighted graphs). Our findings show that the prospective sense of agency may be rooted in the intrinsic functional connectivity patterns of the same regions that are modulated during motor control. Additionally, they suggest that altered inter-network brain connections may shift the individual predisposition to contingent effects conveyed by the contextual sensory feedback, which consequently leads to diversified behavioral responses within the population. To note, the participation coefficients, but not nodal efficiency, were significantly associated with intentional binding. This suggests that the sense of agency is entailed in the modular structure of the brain since participation coefficients are module topography-dependent while nodal efficiency is module topography-independent. Conceivably, stronger inter-network connections mediated by this fronto-parietal subsystem may be necessary for integrating action consequences with ongoing motor behavior to efficiently acquire a feeling of control over actions and their consequences.

Another intriguing finding in our study concerns the relationship between the prospective sense of agency and global modularity, but not global efficiency. Post-hoc analyses further showed that nodes in the primary sensorimotor and middle-insular cortices could be the primary source of this association. Global modularity parameters obtained during a task-free state may be interpreted as describing intrinsic organizational principles that quantify how efficiently a brain is represented by subnetworks, or modules (Gallen and D’Esposito, 2019). Previous studies have reported various associations between whole-brain parameters and behavior, such as between the modular organization and intelligence (Hilger et al., 2017), between inter-modular connections and awareness (Godwin et al., 2015), and between global efficiency and short-term memory skills (Gupta et al., 2018). Expanding on these studies, our findings suggest that brain-behavior associations may exist at multiple hierarchical levels of brain architecture, especially regarding module topography-dependent measures.

It should be noted here that modularity measures may suffer from structural constraints (Betzel et al., 2016). This is a direct consequence of how the modularity itself is estimated: the observed intra-modular connections are compared with the expected intra-modular connections, which are estimated using null models (Newman and Girvan, 2004; Bassett et al., 2011). The constraint in this procedure arises because the weight of the null model is usually assumed to be unitary (γ = 1), essentially constricting the structural resolution of the network (i.e., the size of the modules). In this study, we overcame this limitation by exploring multiple resolution parameters. The results showed that increased values of modularity within medium-high structural resolutions were associated with increased prospective SoA. This interaction was significant with all the regression models that we tested, using both classic and γ-adjusted modularity, and using both binary and weighted graphs. These findings indicate that whole-brain features like modularity may explain high-order mechanisms like contextual predictions elicited by SoA when the brain subsystems (modules) fall between a medium and a small size. Thus, specific resolutions of the modular structure may support specific cognitive functions.

We suggest that the relationship between the sense of agency and global modularity might be explained by the reliance of prospective intentional binding effects on multimodal predictive mechanisms constituting a general principle of brain functioning (Wolpert, 1997; Körding et al., 2007; Clark, 2013). In line with this hypothesis, several researchers have proposed a link between predictive coding and the self (Friston, 2012; Kannape and Blanke, 2012; Apps and Tsakiris, 2014). For instance, an essential feature of the self is the ability to generate probabilistic inferences as efferent copies (Feinberg, 1978; Synofzik et al., 2008; Friston and Kiebel, 2009), which are then compared with sensory perceptions (Wolpert et al., 2001; Apps and Tsakiris, 2014). According to internal prediction models, such comparisons are needed to generate prediction errors (Sato and Yasuda, 2005) that favor the adjustment of the motor response (Bédard and Sanes, 2014). The relevance of such considerations is underlined by the crucial importance of the sense of agency for self-awareness (Gallagher, 2000; Taylor, 2014; Ruvolo, 2015; Haggard, 2017). Further research is needed to directly explore the interplay between global brain properties and predictive coding.

The relation between the sense of agency and the integrative functioning of brain networks may also potentially impact clinical contexts. We argue that a suboptimal prospective sense of agency, commonly observed in neuropathological conditions related to distributed cortical abnormalities such as psychosis (Voss et al., 2010; Moore and Fletcher, 2012; de Bézenac et al., 2018) and in altered awareness (Schooler et al., 2014; Berberian et al., 2017), may arise through two non-exclusive neurophysiological processes. On the one hand, failure of the brain to adapt its structural resolution to the ongoing behavioral context may favor polluted probabilistic estimates of action consequences. On the other hand, inaccurate sensorimotor predictions may result from aberrant intrinsic patterns of nodal functional connectivity in specific brain modules involved in agency tasks. Our report provides a first neurophysiological whole-brain backbone to psychological models of a prediction-based sense of agency that can be evaluated in clinical samples of psychopathology.

Some limitations of this study need to be mentioned. The study focuses on implicit (prospective and retrospective) measures of the sense of agency since explicit judgments of agency may be affected by cognitive biases (Wegner, 2002; Synofzik et al., 2008), implicit beliefs (Desantis et al., 2011; Hoogeveen et al., 2018), and attentional biases (Wen et al., 2016). Accordingly, we argue that implicit behavioral measurements, rather than self-report measures, could be more suitable to start exploring the link between the sense of agency and intrinsic features of the brain. Moreover, this study replicates intentional binding effects from previous investigations (Voss et al., 2010; Di Plinio et al., 2019a), supporting the robustness of the prospective intentional binding effect that reflects the implicit sense of agency (Moore and Haggard, 2008; Moore and Obhi, 2012). This cross-study robustness should also dispel potential concerns about the imbalance of the number of trials across conditions. Some general problems related to studies based on individual differences need to be mentioned, such as low experimental control. To limit such potential confounds, control analyses were performed using the years of sports training and music expertise as covariates (Herting and Nagel, 2013; Kühnis et al., 2014), showing that the effects reported in the present study were independent of these factors. Finally, the sample size may be considered small for individual differences analyses. However, the reliability of our results was demonstrated by applying bootstrapping procedures at the nodal, modular, and global levels. Furthermore, results were confirmed across multiple thresholds (retaining from 10% to 30% of the connections) and using both binary and weighted graphs.

In conclusion, we showed that both whole-brain intra-modular connections and local intrinsic inter-modular connections support the prospective sense of agency. Comparing predictions and intentions with action feedback is an essential feature of human brain function. This process may be favored by a neural substrate showing, on the one hand, a modular architecture represented by highly interconnected, adaptably rewiring subsystems and, on the other hand, efficient exchange of information mediated by motor-control brain systems. The balance of these two components may predispose to more efficient processing of extrinsic self-information, while aberrant network behavior may be a potential marker of increased psychosis-like traits (Di Plinio et al., 2019b). Future studies may consider combining implicit and explicit measures of agency to distinguish between contextual effects and personal biases and their relationships with interindividual differences in brain activity and connectivity.

## Acknowledgments

This work was supported by the BIAL Foundation grant number 195/16 to SE, and by the “Departments of Excellence 2018-2022” initiative of the Italian Ministry of Education, University and Research for the Department of Neuroscience, Imaging and Clinical Sciences (DNISC) of the University of Chieti-Pescara.

